# Identifying factors that contribute to the behavioral effects of psilocybin in preclinical mouse models

**DOI:** 10.64898/2026.07.24.740586

**Authors:** Sixtine Fleury, Luke J. Chang, Katherine M. Nautiyal

## Abstract

Psychedelic drugs offer a potentially promising avenue for novel therapeutics for the treatment of psychiatric conditions, including for mood and anxiety disorders. Emerging clinical studies have prompted increased preclinical research into mechanisms of action. However, post-acute behavioral outcomes in rodents remain inconsistent and difficult to reproduce across laboratories. To address this, we collected a large dataset of behavioral responses to psilocybin in mice (N=693) and used data-driven analyses to identify biological and experimental factors that modulated behavioral readouts. We examined the effects of sex, age, strain, stress protocol, and 5-HT1B signaling on the response to a single high dose of psilocybin measured in acute (head twitch responding and locomotion) or post-acute (sucrose preference, elevated plus maze, novelty-suppressed feeding) behavioral outcomes. We found that psilocybin displays a robust acute behavioral response and a more variable post-acute behavioral profile with a most robust response in sucrose preference. Sex, stress model, and 5-HT1BR activation emerged as key modulators of post-acute antidepressant-like and anxiolytic effects. Random forest classification was narrowly able to predict drug treatment based on post-acute behavioral outcomes and biological and experimental factors. These findings suggest that the heterogeneity seen in the response to psilocybin in mice reflects meaningful biological and experimental variables. Accounting for these factors may enhance translational relevance by identifying the conditions under which therapeutic-like effects are most robust and can support more reproducible models for understanding the neural mechanisms underlying the persisting behavioral effects of psilocybin.

## INTRODUCTION

There has been a surge of interest in psychedelic medicine, especially for psychiatry. Clinical trials have recently shown treatment effects of psychedelics in participants diagnosed with a number of disorders including major depressive, anxiety, post-traumatic stress, and substance use disorders. Among these, psilocybin treatment for depressive and anxiety disorders is among the most prevalent of the published clinical trials (1–5) with multiple studies reporting rapid and sustained symptom improvement lasting weeks to months beyond the acute drug effects (6). Phase 2 trials have demonstrated encouraging signs of safety and efficacy (7) particularly given the substantial unmet need among patients with treatment-resistant forms of depression, for whom existing pharmacotherapies provide limited or no benefit (8). Despite the promising outcomes, clinical trials are conducted in highly controlled settings and are constrained by considerable financial, logistical, and regulatory barriers, limiting scalability and mechanistic inferences. At the same time, growing public interest and rapid clinical translation for novel therapeutic targets have prompted the need to understand the mechanisms through which psilocybin produces durable therapeutic effects.

Preclinical models are critical for understanding the neural mechanisms through which psilocybin has enduring effects on neuropsychiatric phenotypes. However, it has been difficult to establish robust and reproducible post-acute behavioral effects in mice. While acute behavioral and physiological responses to psilocybin are well characterized in rodents (9–11), persisting behavioral effects in standard behavioral pharmacology assays have been inconsistent across studies. For example, contradictory effects of psilocybin have been reported on measures of behavioral despair such as the forced swim test (12–15), with antidepressant-like effects varying considerably depending on the rodent model used (13,16,17). Additional studies have pointed to sex-(18,19) and stress-dependent (20–22) effects of psilocybin, adding further complexity to identifying robust and reproducible post-acute preclinical models. These sources of variability may contribute to discordant results from recent studies examining the role of the 5-HT2AR in the post-acute behavioral response to psilocybin (12,14,23). Critically, even collaborating labs across institutions have had difficulty reproducing post-acute antidepressant-like and anxiolytic effects of psilocybin (24).

The observed variance across labs and assays likely arises from a number of moderating biological and experimental factors (24–26). Commonly used assays of anxiety- and depressive-like behavior in rodents exhibit substantial within- and between-subject variability and a high sensitivity to subtle environmental and procedural differences. Furthermore, the behavioral effects of psilocybin are most likely influenced by multiple interacting biological and experimental factors including dose, timing of assessment, sex, genetic profile and rodent model, stress model, and experimental setup. Thus, detecting and interpreting the post-acute and lasting behavioral effects of psilocybin in preclinical models requires refined experimental approaches that can capture the full multidimensional variability of behavior and neurobiology. Between-subject variability may reflect biologically meaningful differences in susceptibility and treatment response. Identifying factors associated with robust anxiolytic and antidepressant-like response to psilocybin in mice could aid in the improvement of robust preclinical models and provide a critical translational link to clinical applications. Furthermore, the observed variance in post-acute behavioral outcomes across cohorts while optimizing the behavioral assay in our prior work (21), even within our own laboratory conditions, motivated us to examine the biological and experimental sources of this variability within our own dataset.

In the present study, we sought to identify biological and experimental factors that shape the acute and post-acute behavioral effects psilocybin in mice. Using a large, multi-factorial dataset incorporating sex, stress model, strain, age, 5-HT1BR expression, and behavioral outcomes, we assessed psilocybin’s effects on anxiety- and depressive-like phenotypes using complementary statistical modeling and machine learning approaches. We first characterized drug effects on acute behaviors, then evaluated psilocybin’s effects on a shared latent dimension of post-acute behavioral responding before examining each assay individually. Finally, we trained a classifier to discriminate drug assignment across all factors. Our analysis reveals distinct predictors of drug response in acute versus post-acute behaviors, suggesting dissociable mechanisms of action. Sex, stress, and 5-HT1BR signaling emerged as key moderations of post-acute behavioral outcomes. Anxiolytic effects in the elevated plus maze (EPM) and novelty-suppressed feeding test (NSF) were most strongly moderated by stress and 5-HT1BR signaling, while hedonic responding in a gustometer (SPT) showed the most robust antidepressant-like effects overall. Overall, sex emerged as the most robust moderator with females displaying greater sensitivity to psilocybin compared to males.

## MATERIALS AND METHODS

### Animals

Mice were bred in the Dartmouth College vivarium and weaned at postnatal day (PN) 21 into cages of 2–4 same-sex littermates. C57BL/6J mice were obtained from Jackson Laboratory (Strain 00664) to establish a colony. 129S6/SvEv;C57BL/6J mice were bred in the lab from homozygous floxed-tetO1B mice. Mice lacking 5-HT1BR expression and littermate controls were generated by crossing female homozygous floxed-tetO1B mice to male homozygous floxed-tetO1B mice with the βActin-tTS transgene (tetO1B+/+ females crossed to tetO1B+/+::βActin-tTS + males), as previously reported and validated in (27). Mice were maintained on a 12 h light/dark cycle on *ad libitum* food and water until experimental testing. Mice were between the ages of 10-48 weeks old at time of testing. A total of 693 mice (393 males, 300 females) were included in this analysis, comprising N = 128 C57BL/6J mice and N = 565 129S6/SvEv;C57BL/6J mice (N = 224 5-HT1BR knock-outs, N = 364 congenic littermates). Acute behaviors were measured in N = 85 mice (50 males, 35 females). Mice were tested across 15 different cohorts across a three-year span. Of the 565 mixed background mice, data from 371 mice have previously been reported in Fleury & Nautiyal, 2025. See Supplemental Table 1 for numbers of mice run in each behavioral assay stratified by sex, drug condition, genotype, strain, stress, and age. All procedures were approved by the Dartmouth College Institutional Animal Care and Use Committee.

### Drugs and treatments

Psilocybin (Cat # 14041, Cayman Chemical), or saline vehicle, was injected intraperitoneally (i.p.) at the dose of 3mg/kg in C57BL/6J mice and 5mg/kg in tetO1B mice, diluted in saline solution (0.9% NaCl) at a volume of 10ml/kg. The doses were chosen based on our pilot experiments in both strains that induced head twitches and post-acute behavioral changes. 5-HT1B receptors were acutely blocked during drug administration in the C57BL/6J strain by pairing the administration with an i.p injection of 5-HT1B/1D receptor antagonist GR127935 (Cat # 508014; Sigma-Aldrich) at 10mg/kg (in 0.9% saline at 10ml/kg volume) 30 minutes prior to psilocybin or saline vehicle administration, as previously used for pretreatment in behavioral pharmacology studies to target 5-HT1B/D receptors.

#### Chronic corticosterone administration

Corticosterone (CORT) (35 μg/mL; Cat # 2505, Sigma-Aldrich) was dissolved in 0.45% beta-cyclodextrin (Cat # 4767, Sigma-Aldrich) in drinking water, and delivered *ad libitum* in opaque bottles for 4 weeks as described in (21). This paradigm produces a corticosterone dosing of approximately 9.5 mg/kg/day (51). For non-stressed mice, only 0.45% beta-cyclodextrin was dissolved in drinking water and delivered *ad libitum* in opaque bottles for 4 weeks. This was administered for 4 weeks prior to behavioral assay and continued throughout behavioral testing.

#### Chronic behavioral despair model

As an alternative to chronic corticosterone administration, a separate group of male mice were subjected to repeated stress exposures via five forced swim sessions (RFS). Each mouse was placed in a 2-liter beaker containing water (22-25°C) for 10 minutes daily for five consecutive days. Mice remained in their home cages between sessions. Mice were given 48 hours of rest before drug injection.

### Behavioral assays

#### Head Twitch Response and Locomotion

Head twitches and locomotion were analyzed in a subset of mice after the administration of either vehicle or psilocybin as reported in Fleury and Nautiyal, 2025. Mice were placed in an open field immediately after drug administration and recorded for 30 minutes. Head twitches were scored for 15 minutes by three blinded experimenters. Locomotion was quantified using EthoVision for the full 30 minutes. All behavioral testing occurred between the hours of 9am-2pm.

#### Sucrose preference in gustometer

Sucrose licking was tested in a Davis Rig 16-bottle gustometer as reported in detail in Fleury & Nautiyal, 2025. Testing occurred between 9am-2pm. Mice were trained for 5 days in the gustometer chamber where they were exposed to free access of 5% sucrose and 10% sucrose both as a cage and individually for 15 to 30 minutes daily. Following training, mice were food restricted and baseline tested in a 6-bottle paradigm for 30 minutes with access to water, 2% sucrose, 4% sucrose, 6% sucrose, 8% sucrose, and 10% sucrose. Each concentration was randomly presented for a maximum of 120 seconds at a time, and total licks at each tube were recorded using a capacitance-based system. Data were normalized by weight. Mice that had less than 300 licks during the final training session were excluded from the analysis. The 6-bottle paradigm was then repeated 24 hours post drug administration for post-acute behavioral testing. For the purpose of this analysis, sucrose preference score (SPT score) was established from average total licks at the maximum sucrose concentration, 10% sucrose, as a measure of hedonic responding and reward consumption. SPT was measured in 562 mice.

#### Novelty Suppressed Feeding (NSF)

The NSF assay was conducted as reported in detail in Fleury & Nautiyal, 2025. Mice were individually housed 30 minutes prior to NSF testing. Testing occurred between the hours of 3pm-6pm. NSF testing was conducted in a rodent transport container with corn bedding and a single food pellet placed on a white platform in the center of the arena. A lamp was placed above the arena to induce an anxiogenic environment. Mice were placed in the arena for 300 seconds, and latency to eat, defined as the mouse biting the pellet, was recorded over the session. Immediately after a bite, the pellet was removed from the arena. After the session, mice were immediately placed in their home cage with a new food pellet to measure latency to eat within their home cage. Animals that did not bite during the testing session were attributed the maximum value (300 seconds). Animals were placed back in their home cage with food and water ad libitum once all cage mates were tested. NSF was measured in 594 mice.

#### Elevated Plus Maze (EPM)

The EPM was conducted as reported in detail in Fleury & Nautiyal, 2025. Testing occurred between the hours of 3pm-6pm. Mice were placed on the maze (∼100 lux) and allowed to explore into open and closed arms for 6 minutes undisturbed. A camera was placed at the ceiling, and behavior was scored using EthoVision software for duration and number of entries into open arms as a measure of anxiety. For the purpose of this analysis, EPM score was established as the number of entries into open arms. EPM was measured in 554 mice.

### Statistical analysis

Additional details for these statistical analyses are described in the Extended Methods.

#### Principal component analysis

PCA was performed on SPT, EPM, and nsf_risk (sign-flipped so higher = better behavioral performance) variables for animals with complete data (N = 464). Variables were centered and scaled. PCA used singular value decomposition (prcomp in R). Component retention followed Kaiser’s criterion (eigenvalue > 1). Rotation was not applicable in this case given only one component exceeded Kaiser’s criterion. Pairwise Pearson correlations were examined prior to PCA. Loadings, eigenvalues, variance explained, and communalities are reported. PC scores were used in linear regression models to test associations with drug treatment, sex, age, strain, stress, and 5-HT1B expression.

#### Multiple regression

To assess moderators of psilocybin’s acute behavioral effects, we estimated a univariate linear regression model separately for each acute behavioral outcome. Models included drug treatment, age (mean-centered), sex, strain, stress, and 5-HT1BR signaling, along with interaction terms between drug and each covariate to test moderation effects. We report the F-tests using Type III sums of squares for each main effects and interaction. We also fit a reduced model excluding interactions to assess robustness of main drug effects for locomotion.

#### Structural Equation Modeling

To assess which baseline factors (sex, age, strain, stress, and 5-HT1BR signaling) moderate psilocybin’s behavioral effects, we fit a structural equation model using the lavaan package (v.06-20) in R. Sucrose preference (SPT) and elevated plus maze (EPM) performance were specified as outcome variables with freely estimated residual covariance to account for shared variance across domains. Drug (psilocybin vs. saline), age (mean-centered), sex, 5-HT1BR signaling, strain (C57 vs. 129/C57), and stress exposure were included as predictors, with drug x covariate interaction terms to test moderation effects. Continuous predictors were normalized prior to inclusion. Models were estimated using MLR to account for non-normality, with FIML for missing data. Standardized coefficients (β), standard errors, z-values, p-values, and model fit indices (TLI, RMSEA) are reported.

#### Cox proportional hazard regression

NSF latency was analyzed using Cox proportional hazards regression to account for right-censored data. Time to feed was modeled as a function of drug, age, sex, 5-HT1B expression, strain, and stress, with drug × covariate and drug × genotype × stress interaction terms. Non-feeding animals were treated as censored. Hazard ratios and confidence intervals are reported. Proportional hazards assumptions were confirmed via Schoenfeld residuals (global p > 0.05); model concordance was C = 0.671. A continuous NSF-derived anxiety index (nsf_risk) was computed by fitting a null Cox model to NSF latency, extracting deviance residuals, and standardizing them, yielding a z-scored risk index suitable for principal component and machine learning analyses while preserving information from censored observations.

#### Random forest classification

A random forest classifier was trained to assess whether a multivariate signature of psilocybin treatment could be detected from biological and behavioral factors in 5-HT1BR-intact mice (N=288). Predictors included sex, age, strain, stress exposure, SPT, EPM, and NSF risk. Data were partitioned into stratified (70%) and fixed held-out test (30%) sets. The forest was refit 2,000 times on bootstrap resamples of the training data, each time evaluated on the same fixed test set; mean AUC and 95% Cis were derived from this distribution. Variable importance was quantified by mean decrease in accuracy (MDA). Statistical significance was assessed by permutation testing (5,000 iterations): drug labels were shuffled and the model refit on a fresh split each time to generate a null AUC distribution. All analyses used the randomForest and pROC packages in R (seed = 123).

## RESULTS

### Psilocybin causes acute and post-acute behavioral effects in mice

To characterize the acute behavioral effects of psilocybin, we measured the head twitch response and locomotor activity immediately following drug administration (Fig S1). Mice treated with psilocybin demonstrated significantly more head twitching compared to saline-treated mice during the 15 min after administration (Fig. 1B, Welch’s t(34.55) = 11.76, p < 0.0001). This effect was notably modulated by age and strain (Fig. S2). Psilocybin also significantly reduced locomotor activity in the open field compared to saline during the 30 minutes following administration (Fig. 1C, Welch’s t(77.49) = 3.095, p = 0.003).

**Figure 1.**
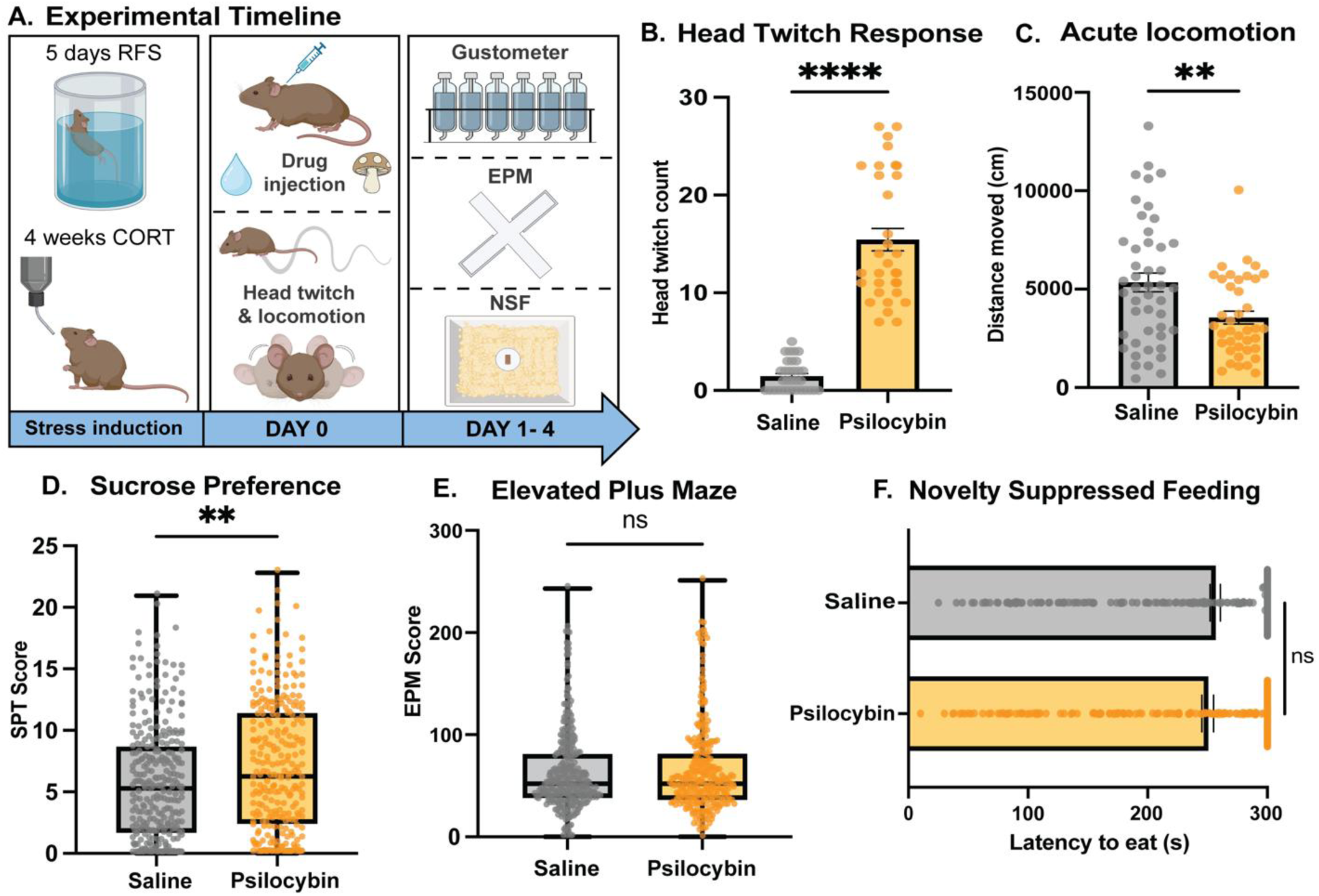
Psilocybin induces robust acute behavioral responses, but variable post-acute behavioral responses. (A) Schematic of the experimental timeline. Mice were subjected to unstressed control conditions or stress induction via repeated forced swim (RFS; 5 days) or chronic corticosterone administration (CORT; 4 weeks). Mice were injected i.p. with saline or psilocybin (5mg/kg), followed immediately by assessment of head-twitch responding for 15 minutes and locomotor activity for 30 minutes. Post-acute behavioral testing was conducted on Days 1-4 and included sucrose preference testing (SPT), elevated plus maze (EPM), and novelty-suppressed feeding (NSF). (B) Acute psilocybin administration induces a robust increase in the head-twitch response in the first 15 minutes after injection. (C) Acute psilocybin administration significantly decreases locomotor activity, as measured by total distance traveled within 30 minutes of injection in an open field. (D) Psilocybin significantly increases SPT scores relative to saline. (E) There were no significant differences observed between saline and psilocybin-treated animals in the EPM. (F) No significant differences were observed between saline- and psilocybin-treated mice in the NSF. Data from individual animals are shown as dots overlaid on bar graphs showing the group mean ± SEM, or boxplots showing median with interquartile range; whiskers indicate range. Statistical differences between treatment groups are indicated as p < 0.05*, p < 0.01**, p < 0.0001 ****, or ns, not significant.

To assess the effects of psilocybin on post-acute behavior, we used assays for anxiety- and depressive-like behavior 1-4 days after injection of saline or psilocybin in 594 mice (see Supplemental Table 1). Mice were tested for anhedonia in the gustometer in a sucrose preference-like test (SPT), and for anxiety-like behavior in elevated plus maze (EPM) and novelty-suppressed feeding (NSF) paradigms. Mice treated with psilocybin showed significantly increased hedonic responding as measured by increased licking for 10% sucrose (Fig. 1D, Welch’s t(536.6) = 3.032, p = 0.003). In the EPM, across all animals, there was no significant difference between the saline-treated and the psilocybin-treated group in exploration into open arms as measured by frequency into open arms (Fig. 1E, Welch’s t(580.6) = 0.4147, p = 0.679). There was also no significant difference in the NSF between saline and psilocybin animals as measured by latency to eat (Fig. 1F, Mann-Whitney U = 36,506 p = 0.290). To directly compare differences in individual variability in behavior between acute and post-acute measures, we also calculated coefficients of variation (CV) for each behavioral outcome by drug group. Locomotion and head twitch responding showed consistently lower variability in psilocybin-treated animals (locomotion CV = 57.8%; HTR CV = 42.3%) compared to psilocybin-treated mice in post-acute measures (SPT CV = 76.3%; EPM CV = 67.3%). Overall, these results suggest that psilocybin results in a robust acute behavioral response, and a more variable post-acute behavioral response. To identify factors contributing to this variability, we performed a series of analyses to determine what covariates contributed to the heterogeneity in post-acute behavioral effects.

### Post-acute behaviors can be represented by a single dimension that is influenced by psilocybin

Turning to the post-acute behavioral effects of psilocybin characterized by high variability, we first examined relationships among the three behavioral measures across animals with complete data for SPT, EPM, and NSF (N = 464). Behavioral measures were weakly correlated (|r| < 0.25), with modest positive associations between SPT and NSF (r = 0.21, p < 0.001), and between SPT and EPM (r = 0.17, p < 0.001), but not between NSF and EPM (r = 0.07, p = 0.13). Similar patterns of correlation were observed within the subsamples of saline-treated animals (SPT-NSF: r = 0.18, p = 0.005; SPT-EPM: r = 0.13, p = 0.039; EPM-NSF: r = 0.11, p = 0.08), and psilocybin-treated animals (SPT-NSF: r = 0.24, p < 0.001; SPT-EPM: r = 0.19, p = 0.005; EPM-NSF: r = 0.03, p = 0.664)(Fig.2A). Direct comparisons of correlation strength between drug groups using Fisher’s r-to-z transformation revealed no significant drug-dependent differences in pairwise correlations (SPT-NSF: p = 0.521, SPT-EPM: p = 0.527, EPM-NSF: p = 0.380), suggesting that psilocybin does not alter the relationship of these behaviors to each other.

**Figure 2.**
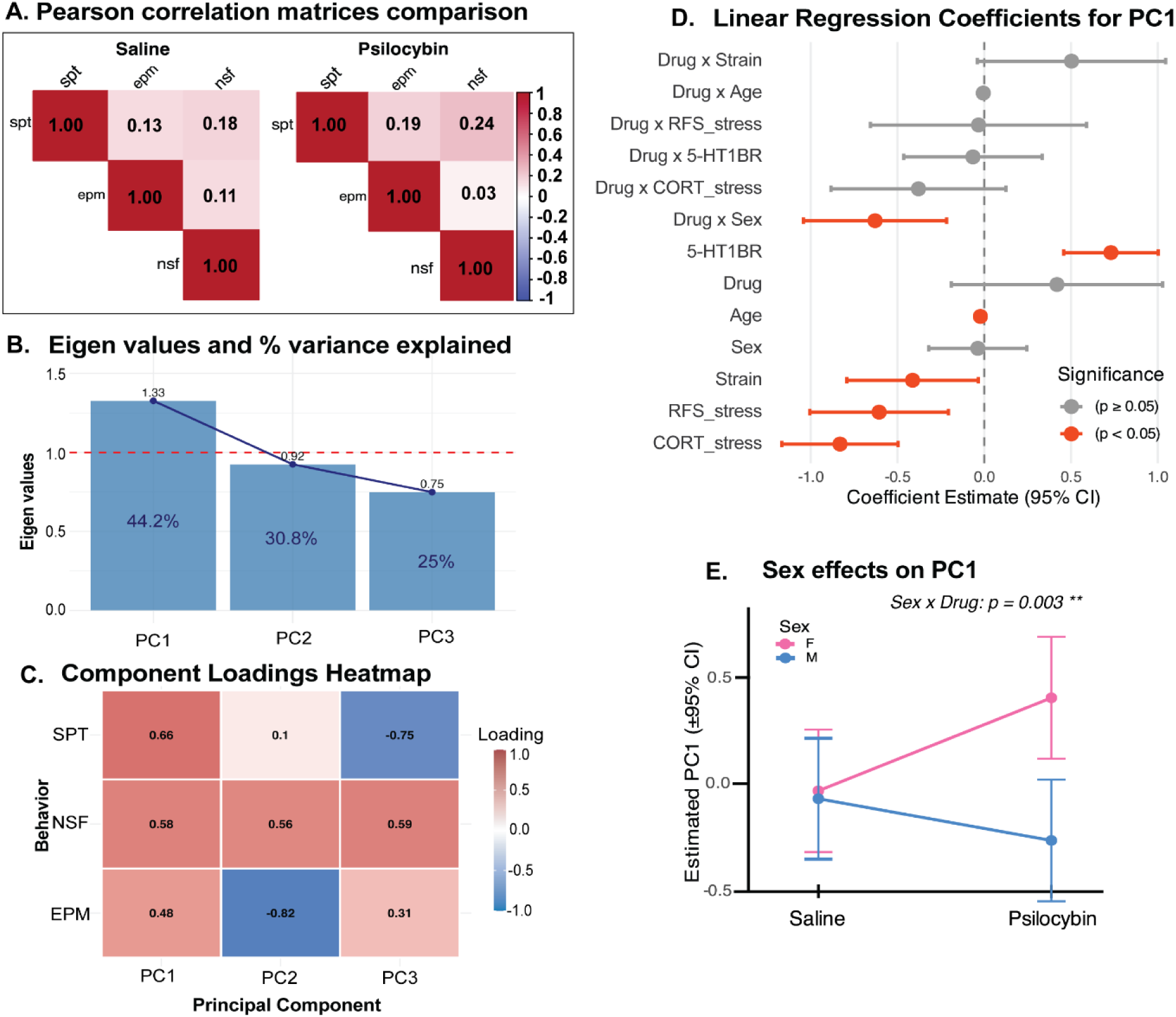
Principal component analysis identifies a shared behavioral axis of affective state which is modulated by psilocybin and baseline variables. (A) Pairwise correlation matrices of behavioral measures following saline (N = 247) or psilocybin (N = 217) treatment. Pearson correlations between SPT, EPM, and NSF show no significant changes following psilocybin administration. (B) Eigen values and percentage of variance explained by each principal component derived from behavioral measures. The first principal component (PC1) accounted for 44.2% of total variance and exceeded the Kaiser criterion (eigenvalue > 1, red dashed line), while PC2 and PC3 accounted for 30.8% and 25.0% of the variance, respectively. (C) Component loading heatmap shows the contribution of each behavioral measure to PC1-PC3. PC1 loads positively on SPT, NSF, and EPM capturing a shared behavioral dimension across assays. PC2 and PC3 captures orthogonal variance with opposing loadings across SPT and EPM. (D) Linear regression model coefficients which predict PC1 scores are shown. Estimated coefficients with 95% confidence intervals for main effects and interaction terms of drug, strain, sex, age, 5-HT1BR, and stress models (RFS, CORT). Significant predictors and interactions (p < 0.05) are shown in red. Stress models, the 129;C57BL/6J strain, and older animals significantly predict lower PC1 scores, while blockade of 5-HT1B predicted higher PC1 scores. (E) Sex-specific effects of psilocybin on PC1. Sex significantly modulates psilocybin’s effect on estimated PC1 scores with psilocybin increasing PC1 scores in females but decreasing PC1 scores in males (p = 0.003).

To assess whether a meaningful behavioral dimension could be identified from the three measures, we performed principal component analysis (PCA) on standardized SPT, EPM, and NSF measures. The first principal component (PC1) had an eigenvalue of 1.33 and explained 44.2% of the total variance, whereas PC2 and PC3 accounted for only 30.8% and 25.0% of variance respectively (Fig. 2B). Following sign alignment of NSF, all three measures loaded positively onto PC1 (SPT = 0.66, NSF = 0.58, EPM = 0.48; Fig. 2C), with higher scores reflecting a positive affective shift including decreased anhedonia, increased exploration into open arms in the EPM, and reduced anxiety-like behavior in the NSF. Linear regression revealed that psilocybin significantly increased PC1 (β = 0.26, p = 0.008; Supp. Table 2), shifting the behavioral dimension towards the positive affective state. We assessed what factors moderated the effects of psilocybin on PC1 by fitting an interaction model including drug x covariate terms (Fig 2D). A significant drug x sex interaction was found (β = −0.63, p = 0.003), with psilocybin increasing PC1 scores in females, but not in males (Fig. 2E). Together, these analyses indicate that there is a small shared component of variance across our three post-acute behavioral measures that may represent a measure of positive affective behavior that is sensitive to psilocybin and modulated by sex.

### Integrated multivariate modeling confirms effects of drug, age, stress, sex, and strain on individual post-acute behavioral measures in mice

While the PCA identified a relatively modest shared dimension of post-acute behavioral responding moderated by psilocybin and sex, the majority of the variance in each measure remained independent, suggesting that each assays captured different facets of overall affective state. We therefore modeled each behavior independently to identify outcome-specific predictors while accounting for the residual covariance between measures. Using structural equation modeling, we found that psilocybin significantly increased hedonic consumption in the SPT (b = 2.352, p = 0.025), while showing only a trend towards increased exploration in the EPM (b = 27.238, p = 0.073; see Supp. Table 3). Several biological factors showed main effects consistent with prior reports, supporting the validity of our approach (28–30) For example, in the SPT, there were significant main effects of age (b = −0.113, p = 0.002), sex (b = −1.422, p = 0.011), and two stress paradigms (CORT: b = −2.374, p < 0.001; RFS: b = −1.973, p = 0.012) such that older animals, males, and stressed mice showed lower sucrose licking scores, suggesting lower hedonic responding. In the EPM, there were significant main effects of sex (b = 9.354, p = 0.035) and 5-HT1BR (b = 11.320, p = 0.009) such that males and 5-HT1BR– mice showed more exploration in the EPM, suggesting lower anxiety. There were also significant main effects of strain (b = −37.429, p < 0.001) and stress models (CORT: b = −30.077, p < 0.001; RFS: b = −32.886, p < 0.001) such that 129;C57BL/6J mice and stressed mice displayed lower EPM scores than C57BL/6J and unstressed mice, respectively. Residual covariance between SPT and EPM remained significant after covariate adjustment (residual covariance estimate = 16.703, p = 0.009), confirming modest shared variance beyond the modeled predictors. Using survival modeling to examine behavior in the NSF, we found that psilocybin did not significantly affect behavior overall, though CORT-stressed mice showed increased latency relative to unstressed mice suggesting increased anxiety-like behavior (HR = −0.80, p = 0.002), consistent with prior reports (31).

### Sex and stress influence the post-acute behavioral responses to psilocybin in mice

Given the large individual differences in these behavioral assays, we explored how biological variables and stress moderate psilocybin’s effects (Fig 3). Sex significantly moderated psilocybin effects on both SPT and EPM (Fig 3A, 3B). In both behaviors, psilocybin had much larger effects in females compared to males (SPT: b = −2.077, p = 0.014) and (EPM: b = −19.2, p = 0.002). There was no statistically significant effect of sex the response to psilocybin in the NSF (drug x sex interaction: HR = 0.85, p = 0.557; Fig. 3C), though quantitively, the effect was in a similar direction, with females showing a greater anxiolytic response. Despite strong main effects of stress on some of these behavioral measures, there were varied interactions with the response to psilocybin. For the SPT, there was no effect of stress on the response to psilocybin (drug x CORT interaction: p = 0.291, drug x RFS interaction: p = 0.679), indicating that psilocybin significantly increased SPT scores in both unstressed and stressed mice (Fig. 3D). On the other hand, in the EPM, stress was a significant moderator of psilocybin’s effects of both stress paradigms on the response to psilocybin, though only trending under RFS stress (CORT: b = −31.493, p = 0.002, RFS: b = −17.131, p = 0.058). However, surprisingly, stressed mice showed an attenuated effect of psilocybin compared to control mice (Fig. 3E). In the NSF, there was no significant effect of either stress paradigm (CORT: HR = 0.052, p = 0.873, RFS: HR = 0.445, p = 0.245) on the response to psilocybin (Fig. 3F). Overall, these interactions highlight important experimental considerations for examining the post-acute behavioral effects of psilocybin.

**Figure 3.**
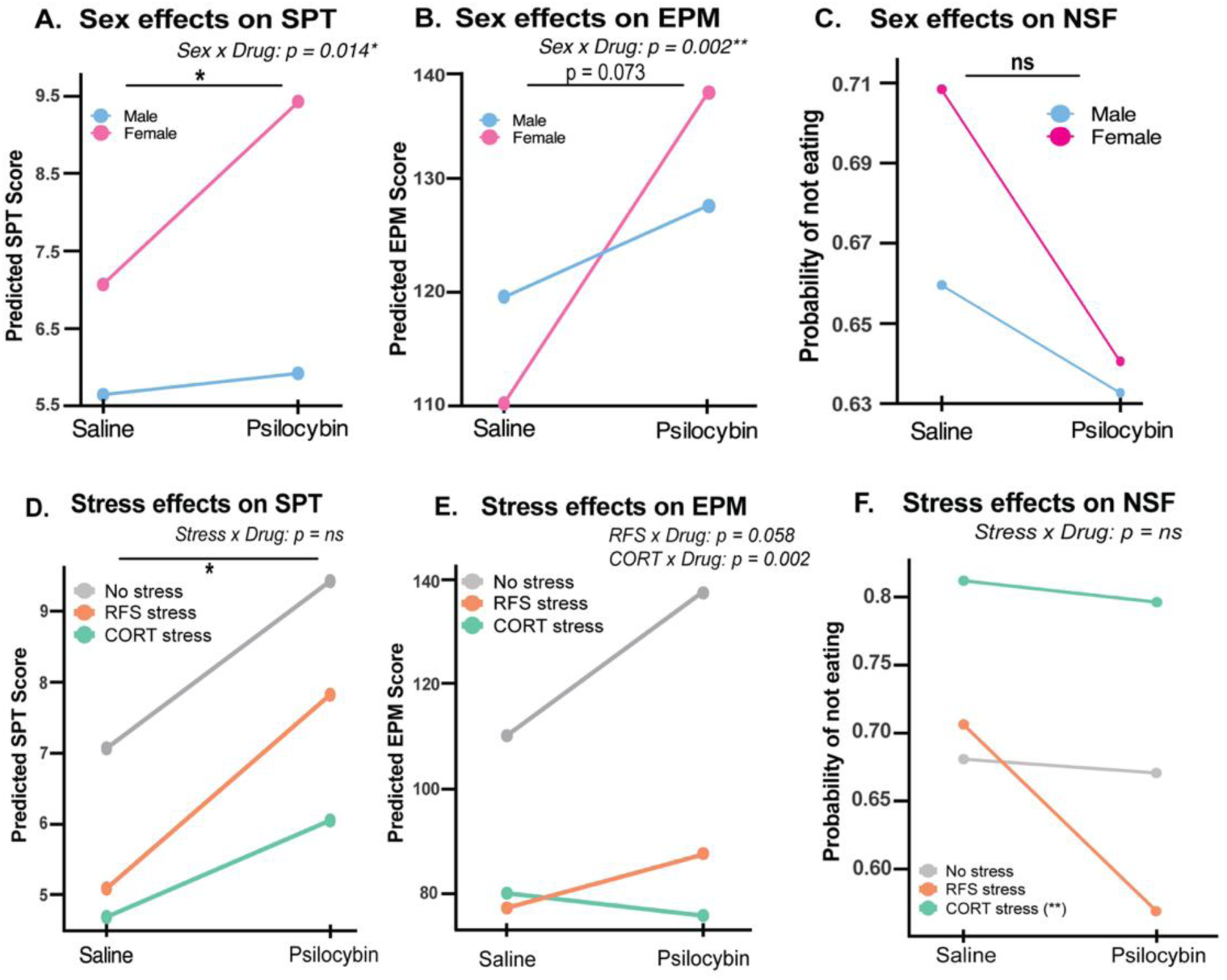
Sex and stress influence the effects of psilocybin in the SPT, EPM, and NSF. (A) Model predicted SPT score shows a significant Sex x Drug interaction (* p < 0.05). Psilocybin increases SPT scores preferentially in females (pink) compared to males (blue). (B) Model predicted EPM scores show a significant Sex x Drug interaction (**p<0.01). Predicted EPM scores indicate a greater psilocybin-induced increase in EPM scores in females relative to males. (C) Model-estimated probability of not eating at the 300-second mark during the NSF task shows a decrease in latency to eat in psilocybin-treated animals preferentially in females compared to males. (D) RFS (orange) and CORT (green) stressed mice display significantly lower SPT scores compared to unstressed mice (gray) (RFS, p = 0.012; CORT, p < 0.0001). Psilocybin significantly increases SPT scores across all stress models, including in unstressed mice. There were no significant differences between the two stress models. (E) CORT and RFS-treated mice show significantly lower EPM scores (p<0.0001; p<0.0001 respectively) compared to unstressed mice. Unstressed mice show a greater increase in EPM exploration following psilocybin compared to RFS stressed mice. CORT stress had a significant effect on psilocybin-induced responding as shown by the decrease in EPM scores in CORT mice compared to an increase in EPM scores in unstressed and RFS mice. (F) There was no significant effect of stress on psilocybin’s response in the NSF at the 300-second mark. Predicted values in Panels A, B, D, and E are derived from a multivariate regression model including age, sex, strain, 5-HT1BR, stress models, drug, and drug x predictor interaction terms. Predicted values in Panel C and F are derived from a Cox proportional hazards regression including age, sex, strain, 5-HT1BR, stress models, drug, and a drug x stress interaction. Error bars indicate 95% confidence intervals. Statistical significance is indicated as p < 0.05*, p<0.01**.

### Multivariate modeling reveals 5-HT1BR as a moderator of post-acute effects of psilocybin

Given that preclinical rodent models are useful for probing mechanistic understanding of the effects of psilocybin on persisting behavioral changes, we examined the ability to detect genetic moderators of these post-acute behavioral effects. Building on our previous findings implicating 5-HT1BR in psilocybin’s antidepressant-like and anxiolytic effects under optimized experimental conditions (21), we now examined three-way interactions in the entire set of mice, focusing on 5-HT1BR mediation of the sex and stress effects (shown in Fig 3). In the EPM, significant drug x stress x 5-HT1BR interactions emerged (Fig. 4A). In mice administered CORT, psilocybin increased anxiety-like behavior in the EPM (read out by decreased EPM scores) only in mice lacking 5-HT1BR signaling – achieved either through pharmacological or genetic manipulations (b = −40.552, p = 0.032). Under RFS, psilocybin also increased anxiety-like behavior in mice lacking 5-HT1BR signaling, but decreased anxiety-like behavior in intact mice (b = −51.084, p = 0.008). These results indicate that psilocybin’s effect on anxiety-related behavior in the EPM is modulated jointly by stress exposure and 5-HT1BR signaling.

**Figure 4.**
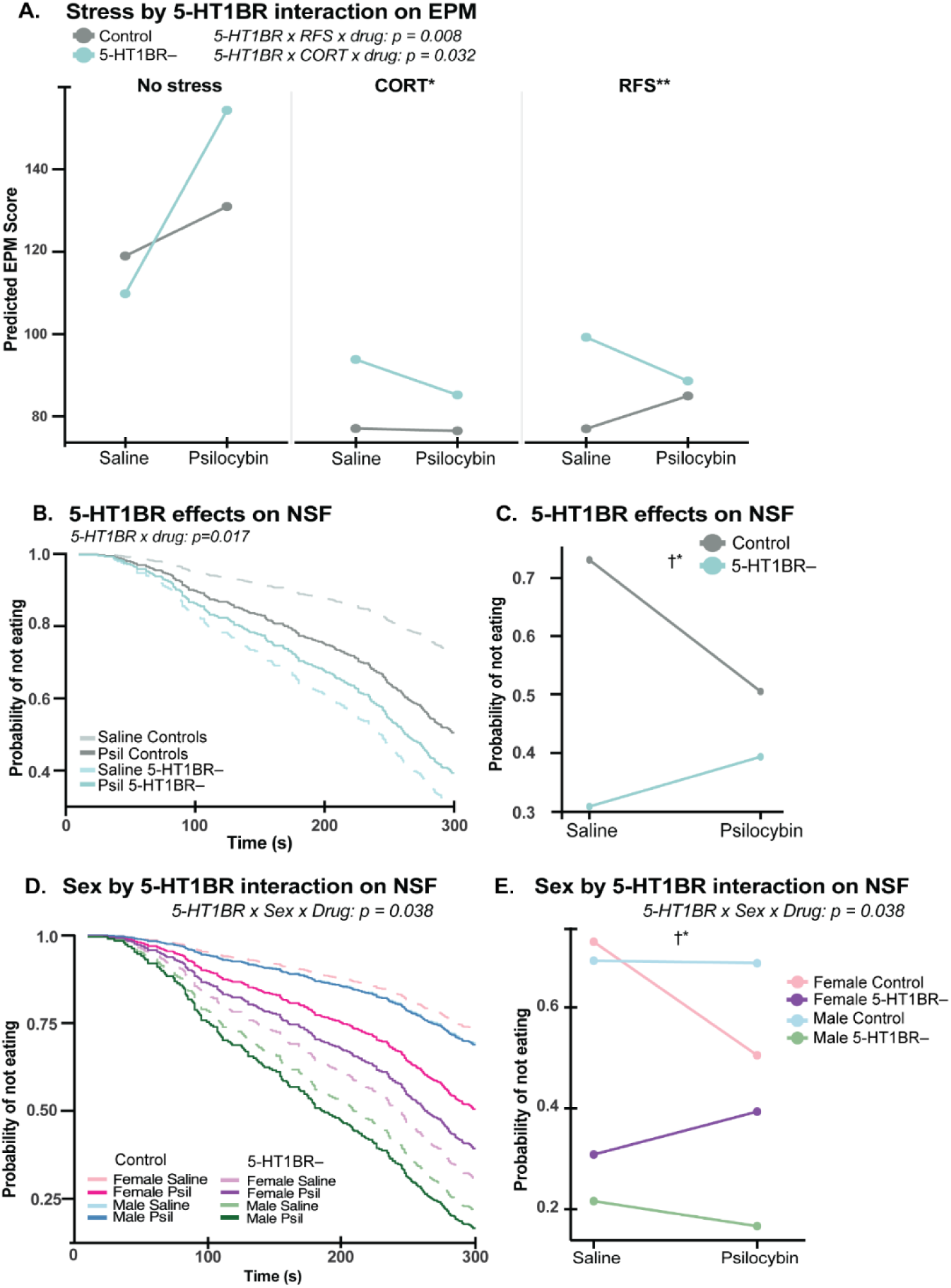
5-HT1BR modulates stress- and sex-dependent effects of psilocybin on anxiety-related behavior. (A) Blockade of 5-HT1BR signaling affects the post-acute effects of psilocybin on EPM behavior differently across stress models. 5-HT1BR– mice (teal) show higher EPM scores overall compared to controls (gray) (p = 0.009). In non-stressed mice, psilocybin increased EPM scores in both control and 5-HT1BR– mice, with a greater effect in 5-HT1BR– mice. Under CORT and RFS stress, psilocybin’s effects were significantly reversed in 5-HT1BR mice. Control mice showed overall no difference in EPM scores under CORT stress, but displayed increased EPM scores under RFS stress. (B)-(C) Blockade of 5-HT1B receptors affect NSF performance following psilocybin administration. Kaplan-Meier survival curves show that 5-HT1BR– mice have an overall higher probability of eating within 300 seconds (p<0.0001). Psilocybin decreases the probability of not eating in control mice, but it has the opposite effect in 5-HT1BR– mice (5-HT1BR x Drug, p = 0.017). (D)-(E) Sex and 5-HT1BR interact to moderate psilocybin’s response in the NSF. A three-way interaction of Drug by Sex by 5-HT1BR reveals significant differences in psilocybin’s response across sex and 5-HT1B blockade (5-HT1BR x Sex x Drug, p = 0.038). Survival curves show that males 5-HT1BR– (green lines) show lower probabilities of not eating compared to control males (blue lines). Psilocybin had no effect in control males (blue), but slightly decreased the probability of not eating in 5-HT1BR– males (green) at the 300-second mark. Female 5-HT1BR– (purple lines) also show lower probabilities of not eating compared to control females (pink lines) at the 300-second mark. Psilocybin had an opposite effect in females compared to males as it increased probabilities of not eating in 5-HT1BR– females (purple), while it decreased it in female controls (pink). Predicated values in panel A are derived from a multivariate regression model including main effects and interaction terms of drug, stress, and 5-HT1BR. Predicated values in panels B through E were derived from using Cox proportional hazards regression including main effects and interaction terms for drug, stress, sex, and 5-HT1BR. Lines represent model-estimated effects. Significance is denoted as p < 0.05*, p < 0.01**. Interaction is denoted by †.

In the NSF, our analysis showed that 5-HT1BR signaling also influenced the 5-HT1BR response to psilocybin (drug × 5-HT1BR: HR = 0.37, p = 0.017), as we previously reported (21). Interestingly while psilocybin decreases anxiety-like behavior in controls (HR = 2.12, p=0.015), psilocybin has no significant effects on anxiety in the absence of 5-HT1BR signaling (HR = 1.004, p=0.85; Fig. 4B, C). A significant three-way drug × 5-HT1BR × sex interaction was also detected (HR = 3.15, p = 0.038; Fig. 4D), with psilocybin reducing anxiety-like behavior in intact females but not males, and this pattern reversing in females lacking 5-HT1BR signaling (Fig. 4E; Supp. Table 4). Together, these findings indicate that sex-specific modulation of psilocybin’s behavioral effects is partly mediated by 5-HT1BR signaling and confirm our previous findings that 5-HT1BR signaling is a critical modulator of psilocybin’s post-acute behavioral effects, gating both the direction and magnitude of its anxiolytic response in a sex- and stress-dependent manner. Overall, these analyses also point to the preclinical post-acute behavioral effects being amenable to genetic and circuit dissection to understand the mechanisms through which psilocybin causes persisting behavioral effects.

Given the variability in psilocybin’s behavioral effects across assays and moderators, we sought to examine if we could predict drug treatment based on biological factors and behavioral outcomes. We did this by fitting a classifier to predict saline versus psilocybin treatment, but restricted analyses to intact mice (N = 288) given the heterogeneity in the 5-HT1BR-related responses. Despite being unable to accurately predict treatment significantly above chance using a random forest classifier trained on biological and behavioral predictors (Fig. 5; mean AUC of 0.603 [95% CI: 0.494-0.689] across 2,000 bootstrap iterations, with a permutation p-value of 0.1 from 5,000 iterations), classification performance did trend towards correct prediction compared to chance control. These results highlight an existing signature of psilocybin, but the effect at population level remains modest.

**Figure 5.**
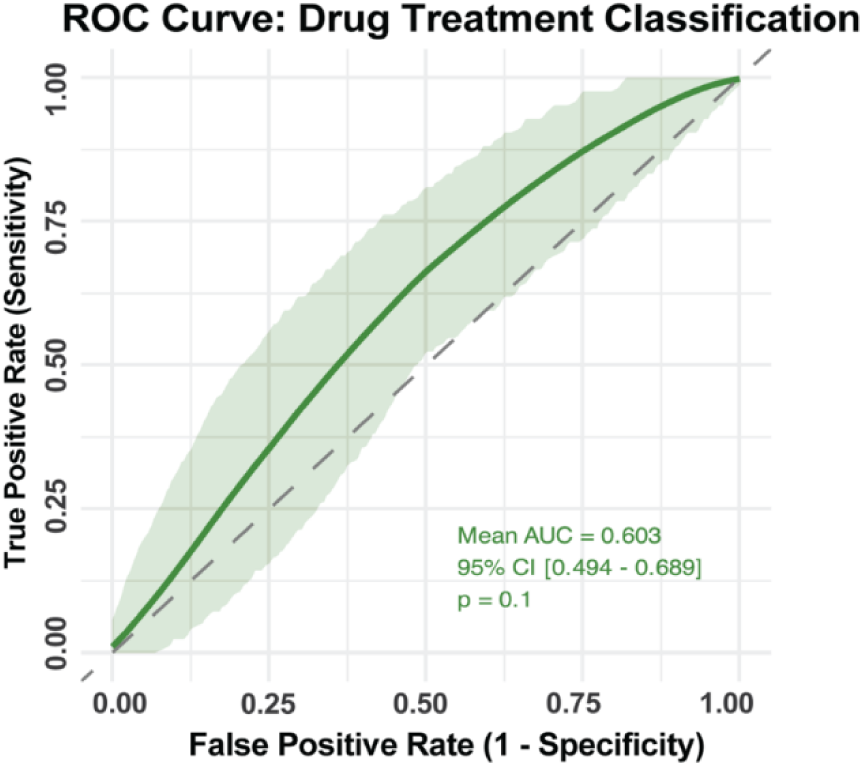
Random forest classification reveals modest prediction of psilocybin treatment. Mean ROC curve across 2,000 bootstrap iterations for a random forest classifier trained to predict drug treatment assignment (psilocybin vs. saline) in 5-HT1BR-intact control mice (N = 288). Predictors included sex, age, strain, stress exposure, and post-acute behavioral outcomes (SPT, EPM, NSF risk). The green line represents the mean ROC curve and the shaded band represents the 95% confidence interval across bootstrap resamples. The dashed diagonal represents chance performance (AUC = 0.5).

## DISCUSSION

By analyzing data from a large set of mice, we were able to identify significant biological and experimental factors that influence the variability of the post-acute behavioral effects of psilocybin. Using a large number of mice, we were only able to find a significant effect of psilocybin in one of three post-acute measures tested (i.e., increased hedonic behavior measured in the SPT). Results from the EPM and NSF showed high variability within treatment groups reflecting low precision and therefore difficult to detect any true effects of psilocybin. Using multivariate analysis, we were able to consider interactions of psilocybin’s effects with biological and environmental variables across this large data set of heterogenous mice. This analysis revealed that the sex and stress exposure of mice reliably influenced the post-acute behavioral response to psilocybin, and that given this information, the SPT and EPM assays are most informative for predicting psilocybin treatment versus saline. We also show that our modeling approach allows for mechanistic investigation, confirming our previous work that 5-HT1BR mediates the post-acute behavioral response in a large heterogenous mouse sample. Overall, our work points to parameters which could be exploited in future studies to increase the robustness and reproducibility of preclinical research investigating the persisting behavioral effects of psilocybin.

Our findings directly address a burgeoning area of work on preclinical models for psilocybin’s behavioral effects, and the associated controversy. A recent report from a multi-institution group highlights the variability of behavioral effects in mice, showing relatively small effects sizes of psilocybin treatment that is dwarfed by between-lab variability (Lu et al, bioRXiv, 2024). While their approach was to minimize sources of variability, our approach was designed to examine these orthogonal sources of variability. Consistent with Lu et al’s interpretation, our multivariate classification analyses achieved only modest discrimination between psilocybin- and saline-treated mice indicating that while psilocybin induces a detectable behavioral profile, the magnitude of this effect is limited, and the effects are subtle at a population level. However, while Lu et al., 2024 reported no significant post-acute behavioral responses to psilocybin across labs, we were able to detect significant behavioral responses especially after controlling for moderating factors such as sex and stressor exposure. Even without accounting for moderating biological and environmental factors, we found a significant effect of psilocybin on post-acute hedonic responding, an antidepressant-like measure. However, this assay included 562 mice which is an impractical target number for future preclinical studies investigating the neurobiological mechanisms of the antidepressant effects of psilocybin. While we agree that psilocybin’s behavioral effects in mice can be relatively modest and difficult to detect, our analysis provides avenues to refine traditional paradigms. By constraining and/or accounting for variables, experiments with more reasonable sample sizes may yield larger effects studies amenable to studying the neurobiological mechanisms underlying the behavioral effects. We conclude that biological variables like sex, stress exposure, and age are not sources of noise but potentially biologically relevant factors that influence the response to psilocybin.

Behavioral effects of psilocybin in mice are interestingly distinguishable from those seen following serotonin reuptake inhibitor (SSRI) administration. One source of difference seems to be related to stress-induction paradigms to promote a depressive-like state that can be “rescued” with SSRI treatment (32–34). In contrast, ours and others’ studies show that stress may diminish antidepressant-like effects of psilocybin (35). Specifically, our results show that psilocybin induces decreases in depressive and anxiety phenotypes from baseline, and stress-induction paradigms reduce the antidepressant and anxiolytic effects of psilocybin. This may actually be a translationally-valid observation, given that psilocybin produces measurable effects on mood and cognition in healthy controls (36), where as an underlying pathophysiology may be required to see behavioral effects of SSRIs (37). Mechanistically, this distinction is also consistent with the divergent ways that SSRIs and psilocybin engage homeostatic circuits related to HPA axis negative feedback seen following SSRIs (38,39), but not psilocybin (40,41). Therefore, it is possible that stress models may provide a better background for studying the effect of SSRIs on behavior because the mechanisms that SSRIs engage overlap more with the stress circuitry. On the other hand psilocybin triggers rapid global reorganization by increasing function integration across brain networks and reducing modularity within the default mode network (42) and promoting rapid and sustained structural remodeling and dendritic spine growth (43,44), tangential to the neural circuits involved in stress. Overall, we propose that the different preclinical models required to see behavioral effects of SSRI or psilocybin treatment suggest a fundamental distinction in circuit-level recruitment.

A primary finding of our analysis is that sex influences the behavioral response to psilocybin. Sex emerged as a significant moderator across all post-acute outcomes from PCA and multivariate regression analyses, with females responding more robustly to psilocybin compared to males. Sex-specific effects of psilocybin have been reported in a number of recent preclinical rodent studies, though the direction of the effect varies based on the behavioral phenotype and/or interactions with other environmental and biological factors. For example, psilocybin reduced opioid preference and withdrawal in males but not in females (19). Psilocybin also increased stimulus specific central amygdala reactivity in females and not males, but it increased neural activity in the paraventricular nucleus of the hypothalamus (PVN) to an aversive air-puff stimulus in males and not females (18,45). Additionally, psilocybin administration reduced ethanol consumption and preference in male mice, but not females (46). Differential effects of sex have also been observed in invertebrate models where psilocybin significantly reduced immobility in the forced swim test in male, but not female flies (47). This suggests that sex differences observed in our results and other rodent research may reflect conserved neurobiological sex differences in the post-acute responses to psilocybin, rather than assay or species-specific artifacts. Interestingly, sex differences have not yet been reported from human trials. It is unclear if this represents a true lack of sex difference in humans, or if trials are not statistically powered or experimentally designed to detect sex differences. Composite scores from self-report questionnaires and clinical ratings may not capture sex-specific changes in the constellation of phenotypic changes which are generally stratified in preclinical studies. Additionally, estrus/menstrual phase could be tracked to help identify effects of hormonal status on behavioral responses. Whether the consistent sex differences seen in preclinical trials indicate a lack of translational relevance or point to detection limitations in human trials, remains an open question.

Overall, we demonstrate that psilocybin induces robust acute responses and more variable post-acute affective changes that are strongly shaped by rodent demographic and experimental factors. Sex and stress significantly moderate behavioral outcomes, emphasizing the need to consider these variables when selecting protocols, and in other aspects of preclinical study design and analyses. Incorporating biological heterogeneity writ large into our preclinical experimental design could improve mechanistic inference and help bridge preclinical behavioral phenotypes with patient stratification approaches in precision psychiatry (57). Future studies can benefit from integrating multivariate behavioral phenotyping with circuit-level measurements to identify neural signatures associated with distinct behavioral response profiles. Specifically, stratifying behavioral responses by these variables could improve sensitivity to detect treatment effects and better reflect variability observed in clinical trials. Ultimately, refining preclinical models to capture more robust behavioral responses will help advance our mechanistic understanding of the persisting behavioral changes to support more precise translation to human therapeutic contexts.

## Acknowledgments

The authors would like to thank Dr. Melissa Herman for helpful discussions.

## Author Contributions

SF and KMN conceived of the study; SF collected data; SF and LJC analyzed data; SF wrote the initial draft of the paper; SF, KMN, and LJC edited the final draft of the paper

## Funding

Dartmouth College Institutional Funds; International Center for Responsible Gaming

## Competing Interests

The Authors have nothing to disclose.

## Notes

### Competing Interest Statement

The authors have declared no competing interest.

## REFERENCES

1. Carhart-Harris RL, Bolstridge M, Rucker J, Day CMJ, Erritzoe D, Kaelen M, et al. Psilocybin with psychological support for treatment-resistant depression: an open-label feasibility study. Lancet Psychiatry. 2016 Jul 1;3(7):619–27. doi:10.1016/S2215-0366(16)30065-7

2. Carhart-Harris RL, Bolstridge M, Day CMJ, Rucker J, Watts R, Erritzoe DE, et al. Psilocybin with psychological support for treatment-resistant depression: six-month follow-up. Psychopharmacology (Berl). 2018 Feb 1;235(2):399–408. doi:10.1007/s00213-017-4771-x

3. Davis AK, Barrett FS, May DG, Cosimano MP, Sepeda ND, Johnson MW, et al. Effects of Psilocybin-Assisted Therapy on Major Depressive Disorder: A Randomized Clinical Trial. JAMA Psychiatry. 2021 May 1;78(5):481. doi:10.1001/jamapsychiatry.2020.3285

4. Grob CS, Danforth AL, Chopra GS, Hagerty M, McKay CR, Halberstadt AL, et al. Pilot Study of Psilocybin Treatment for Anxiety in Patients With Advanced-Stage Cancer. Arch Gen Psychiatry. 2011 Jan 3;68(1):71–8. doi:10.1001/archgenpsychiatry.2010.116

5. Ross S, Bossis A, Guss J, Agin-Liebes G, Malone T, Cohen B, et al. Rapid and sustained symptom reduction following psilocybin treatment for anxiety and depression in patients with life-threatening cancer: a randomized controlled trial. J Psychopharmacol (Oxf). 2016 Dec 1;30(12):1165–80. doi:10.1177/0269881116675512

6. Goodwin GM, Aaronson ST, Alvarez O, Arden PC, Baker A, Bennett JC, et al. Single-Dose Psilocybin for a Treatment-Resistant Episode of Major Depression. N Engl J Med. 2022 Nov 3;387(18):1637–48. doi:10.1056/NEJMoa2206443

7. Li LJ, Mo Y, Shi ZM, Huang XB, Ning YP, Wu HW, et al. Psilocybin for major depressive disorder: a systematic review of randomized controlled studies. Front Psychiatry. 2024 Sep 23;15. doi:10.3389/fpsyt.2024.1416420

8. Gaynes BN, Warden D, Trivedi MH, Wisniewski SR, Fava M, Rush AJ. What Did STAR∗D Teach Us? Results From a Large-Scale, Practical, Clinical Trial for Patients With Depression. Psychiatr Serv. 2009;60(11).

9. Madsen MK, Fisher PM, Burmester D, Dyssegaard A, Stenbæk DS, Kristiansen S, et al. Psychedelic effects of psilocybin correlate with serotonin 2A receptor occupancy and plasma psilocin levels. Neuropsychopharmacology. 2019 Jun;44(7):1328–34. doi:10.1038/s41386-019-0324-9

10. Roseman L, Nutt DJ, Carhart-Harris RL. Quality of Acute Psychedelic Experience Predicts Therapeutic Efficacy of Psilocybin for Treatment-Resistant Depression. Front Pharmacol. 2018 Jan 17;8. doi:10.3389/fphar.2017.00974

11. Yaden DB, Griffiths RR. The Subjective Effects of Psychedelics Are Necessary for Their Enduring Therapeutic Effects. ACS Pharmacol Transl Sci. 2021 Apr 9;4(2):568–72. doi:10.1021/acsptsci.0c00194

12. Hesselgrave N, Troppoli TA, Wulff AB, Cole AB, Thompson SM. Harnessing psilocybin: antidepressant-like behavioral and synaptic actions of psilocybin are independent of 5-HT2R activation in mice. Proc Natl Acad Sci. 2021 Apr 27;118(17):e2022489118. doi:10.1073/pnas.2022489118

13. Hibicke M, Landry AN, Kramer HM, Talman ZK, Nichols CD. Psychedelics, but Not Ketamine, Produce Persistent Antidepressant-like Effects in a Rodent Experimental System for the Study of Depression. ACS Chem Neurosci. 2020 Mar 18;11(6):864–71. doi:10.1021/acschemneuro.9b00493

14. Sekssaoui M, Bockaert J, Marin P, Bécamel C. Antidepressant-like effects of psychedelics in a chronic despair mouse model: is the 5-HT2A receptor the unique player? Neuropsychopharmacology. 2024 Jan 11;1–10. doi:10.1038/s41386-024-01794-6

15. Takaba R, Ibi D, Yoshida K, Hosomi E, Kawase R, Kitagawa H, et al. Ethopharmacological evaluation of antidepressant-like effect of serotonergic psychedelics in C57BL/6J male mice. Naunyn Schmiedebergs Arch Pharmacol. 2024 May 1;397(5):3019–35. doi:10.1007/s00210-023-02778-x

16. Jefsen O, Højgaard K, Christiansen SL, Elfving B, Nutt DJ, Wegener G, et al. Psilocybin lacks antidepressant-like effect in the Flinders Sensitive Line rat. Acta Neuropsychiatr. 2019 Aug;31(04):213–9. doi:10.1017/neu.2019.15

17. Kolasa M, Nikiforuk A, Korlatowicz A, Solich J, Potasiewicz A, Dziedzicka-Wasylewska M, et al. Unraveling psilocybin’s therapeutic potential: behavioral and neuroplasticity insights in Wistar-Kyoto and Wistar male rat models of treatment-resistant depression. Psychopharmacology (Berl). 2025 Jul 1;242(7):1607–25. doi:10.1007/s00213-024-06644-3

18. Effinger DP, Quadir SG, Ramage MC, Cone MG, Herman MA. Sex-specific effects of psychedelic drug exposure on central amygdala reactivity and behavioral responding. Transl Psychiatry. 2023 Apr 8;13(1):1–15. doi:10.1038/s41398-023-02414-5

19. Jaster AM, Hadlock TM, Buzzi B, Maltman JL, Silva GM, Saha S, et al. Sex-specific role of the 5-HT2A receptor in psilocybin-induced extinction of opioid reward. Nat Commun. 2025 Nov 20;16(1):10206. doi:10.1038/s41467-025-64887-w

20. Erkizia-Santamaría I, Horrillo I, Martínez-Álvarez N, Pérez-Martínez D, Rivero G, Erdozain AM, et al. Evaluation of behavioural and neurochemical effects of psilocybin in mice subjected to chronic unpredictable mild stress. Transl Psychiatry. 2025 Jun 14;15:201. doi:10.1038/s41398-025-03421-4 PubMed PMID: 40517150; PubMed Central PMCID: PMC12167372.

21. Fleury S, Nautiyal KM. The serotonin 1B receptor is required for some of the behavioral effects of psilocybin in mice. Mol Psychiatry. 2025 Dec 20;1–13. doi:10.1038/s41380-025-03387-1

22. Wang Z, Robbins B, Zhuang R, van Bruggen R, Sandini T, Li XM, et al. Psilocybin mitigates behavioral despair and cognitive impairment in treatment-resistant depression model using wistar kyoto rats. Sci Rep. 2025 May 26;15(1):18432. doi:10.1038/s41598-025-03383-z

23. Cameron LP, Patel SD, Vargas MV, Barragan EV, Saeger HN, Warren HT, et al. 5-HT2ARs Mediate Therapeutic Behavioral Effects of Psychedelic Tryptamines. ACS Chem Neurosci. 2023 Feb 1;14(3):351–8. doi:10.1021/acschemneuro.2c00718

24. Lu OD, White K, Raymond K, Liu C, Klein AS, Green N, et al. A multi-institutional investigation of psilocybin’s effects on mouse behavior.

25. Erkizia-Santamaría I, Horrillo I, Meana JJ, Ortega JE. Clinical and preclinical evidence of psilocybin as antidepressant. A narrative review. Prog Neuropsychopharmacol Biol Psychiatry. 2025 Jan 10;136:111249. doi:10.1016/j.pnpbp.2025.111249

26. Halberstadt AL, Geyer MA. Multiple receptors contribute to the behavioral effects of indoleamine hallucinogens. Neuropharmacology. 2011 Sep 1;Serotonin: The New Wave61(3):364–81. doi:10.1016/j.neuropharm.2011.01.017

27. Nautiyal KM, Tanaka KF, Barr MM, Tritschler L, Le Dantec Y, David DJ, et al. Distinct Circuits Underlie the Effects of 5-HT1B Receptors on Aggression and Impulsivity. Neuron. 2015 May;86(3):813–26. doi:10.1016/j.neuron.2015.03.041

28. Bertholomey ML, Nagarajan V, Smith DM, Torregrossa MM. Sex- and age-dependent effects of chronic corticosterone exposure on depressive-like, anxiety-like, and fear-related behavior: Role of amygdala glutamate receptors in the rat. Front Behav Neurosci. 2022 Sep 23;16:950000. doi:10.3389/fnbeh.2022.950000 PubMed PMID: 36212195; PubMed Central PMCID: PMC9537815.

29. Xia J, Wang H, Zhang C, Liu B, Li Y, Li K, et al. The comparison of sex differences in depression-like behaviors and neuroinflammatory changes in a rat model of depression induced by chronic stress. Front Behav Neurosci. 2023 Jan 4;16:1059594. doi:10.3389/fnbeh.2022.1059594 PubMed PMID: 36703721; PubMed Central PMCID: PMC9872650.

30. Bowman R, Frankfurt M, Luine V. Sex differences in anxiety and depression: insights from adult rodent models of chronic stress and neural plasticity. Front Behav Neurosci. 2025 May 14;19:1591973. doi:10.3389/fnbeh.2025.1591973

31. Nautiyal KM, Tritschler L, Ahmari SE, David DJ, Gardier AM, Hen R. A Lack of Serotonin 1B Autoreceptors Results in Decreased Anxiety and Depression-Related Behaviors. Neuropsychopharmacology. 2016 Nov;41(12):2941–50. doi:10.1038/npp.2016.109

32. David DJ, Samuels BA, Rainer Q, Wang JW, Marsteller D, Mendez I, et al. Neurogenesis-Dependent and -Independent Effects of Fluoxetine in an Animal Model of Anxiety/Depression. Neuron. 2009 May 28;62(4):479–93. doi:10.1016/j.neuron.2009.04.017 PubMed PMID: 19477151.

33. Ratajczak P, Martyński J, Zięba JK, Świło K, Kopciuch D, Paczkowska A, et al. Comparative Efficacy of Animal Depression Models and Antidepressant Treatment: A Systematic Review and Meta-Analysis. Pharmaceutics. 2024 Aug 29;16(9):1144. doi:10.3390/pharmaceutics16091144

34. Vialou V, Robison AJ, LaPlant QC, Covington HE, Dietz DM, Ohnishi YN, et al. ΔFosB in brain reward circuits mediates resilience to stress and antidepressant responses. Nat Neurosci. 2010 Jun;13(6):745–52. doi:10.1038/nn.2551 PubMed PMID: 20473292; PubMed Central PMCID: PMC2895556.

35. Jones NT, Zahid Z, Grady SM, Sultan ZW, Zheng Z, Banks MI, et al. Delayed Anxiolytic-Like Effects of Psilocybin in Male Mice Are Supported by Acute Glucocorticoid Release [preprint] [Internet]. Animal Behavior and Cognition; 2020 Aug [cited 2023 Jan 30]. Available from: http://biorxiv.org/lookup/doi/10.1101/2020.08.12.248229 doi:10.1101/2020.08.12.248229

36. Barrett FS, Doss MK, Sepeda ND, Pekar JJ, Griffiths RR. Emotions and brain function are altered up to one month after a single high dose of psilocybin. Sci Rep. 2020 Feb 10;10(1):2214. doi:10.1038/s41598-020-59282-y

37. Becker AM, Holze F, Grandinetti T, Klaiber A, Toedtli VE, Kolaczynska KE, et al. Acute Effects of Psilocybin After Escitalopram or Placebo Pretreatment in a Randomized, Double-Blind, Placebo-Controlled, Crossover Study in Healthy Subjects. Clin Pharmacol Ther. 2022;111(4):886–95. doi:10.1002/cpt.2487

38. Doss MK, Madden MB, Gaddis A, Nebel MB, Griffiths RR, Mathur BN, et al. Models of psychedelic drug action: modulation of cortical-subcortical circuits. Brain. 2022 Feb 1;145(2):441–56. doi:10.1093/brain/awab406

39. Tafet GE, Nemeroff CB. Pharmacological Treatment of Anxiety Disorders: The Role of the HPA Axis. Front Psychiatry. 2020 May 15;11. doi:10.3389/fpsyt.2020.00443

40. Mason NL, Szabo A, Kuypers KPC, Mallaroni PA, de la Torre Fornell R, Reckweg JT, et al. Psilocybin induces acute and persisting alterations in immune status in healthy volunteers: An experimental, placebo-controlled study. Brain Behav Immun. 2023 Nov 1;114:299–310. doi:10.1016/j.bbi.2023.09.004

41. Wang *Zitong, Zhang Y, Li XM. PSILOCYBIN MITIGATES BEHAVIORAL DESPAIR AND COGNITIVE RECOGNITION IMPAIRMENTS BY REGULATING THE HYPOTHALAMIC-PITUITARY-ADRENAL (HPA) AXIS VIA THE BRAIN-DERIVED NEUROTROPHIC FACTOR (BDNF) SIGNALING PATHWAY MEDIATED BY THE ENDOCANNABINOID SYSTEM (ECS). Int J Neuropsychopharmacol. 2025 Feb 1;28(Supplement_1):i77–8. doi:10.1093/ijnp/pyae059.134

42. Daws RE, Timmermann C, Giribaldi B, Sexton JD, Wall MB, Erritzoe D, et al. Increased global integration in the brain after psilocybin therapy for depression. Nat Med. 2022 Apr;28(4):844–51. doi:10.1038/s41591-022-01744-z

43. Calder AE, Hasler G. Towards an understanding of psychedelic-induced neuroplasticity. Neuropsychopharmacology. 2023 Jan;48(1):104–12. doi:10.1038/s41386-022-01389-z

44. Shao LX, Liao C, Gregg I, Davoudian PA, Savalia NK, Delagarza K, et al. Psilocybin induces rapid and persistent growth of dendritic spines in frontal cortex in vivo. Neuron. 2021 Aug 18;109(16):2535–2544.e4. doi:10.1016/j.neuron.2021.06.008

45. Effinger DP, Hoffman JL, Mott SE, Magee SN, Quadir SG, Rollison CS, et al. Increased reactivity of the paraventricular nucleus of the hypothalamus and decreased threat responding in male rats following psilocin administration. Nat Commun. 2024 Jun 22;15(1):1–15. doi:10.1038/s41467-024-49741-9

46. Alper K, Cange J, Sah R, Schreiber-Gregory D, Sershen H, Vinod KY. Psilocybin sex-dependently reduces alcohol consumption in C57BL/6J mice. Front Pharmacol. 2023 Jan 4;13:1074633. doi:10.3389/fphar.2022.1074633 PubMed PMID: 36686713; PubMed Central PMCID: PMC9846572.

47. Hibicke M, Nichols CD. Validation of the forced swim test in Drosophila, and its use to demonstrate psilocybin has long-lasting antidepressant-like effects in flies. Sci Rep. 2022 Jun 15;12(1):10019. doi:10.1038/s41598-022-14165-2

48. Karrer TM, McLaughlin CL, Guaglianone CP, Samanez-Larkin GR. Reduced serotonin receptors and transporters in normal aging adults: a meta-analysis of PET and SPECT imaging studies. Neurobiol Aging. 2019 Aug;80:1–10. doi:10.1016/j.neurobiolaging.2019.03.021 PubMed PMID: 31055162; PubMed Central PMCID: PMC6679764.

49. Lambe EK, Fillman SG, Webster MJ, Weickert CS. Serotonin Receptor Expression in Human Prefrontal Cortex: Balancing Excitation and Inhibition across Postnatal Development. PLOS ONE. 2011 Jul 29;6(7):e22799. doi:10.1371/journal.pone.0022799

50. Meltzer CC, Smith G, DeKosky ST, Pollock BG, Mathis CA, Moore RY, et al. Serotonin in aging, late-life depression, and Alzheimer’s disease: the emerging role of functional imaging. Neuropsychopharmacol Off Publ Am Coll Neuropsychopharmacol. 1998 Jun;18(6):407–30. doi:10.1016/S0893-133X(97)00194-2 PubMed PMID: 9571651.

51. Sheline YI, Mintun MA, Moerlein SM, Snyder AZ. Greater Loss of 5-HT2A Receptors in Midlife Than in Late Life. Am J Psychiatry. 2002 Mar 1;159(3):430–5. doi:10.1176/appi.ajp.159.3.430

52. Cummins BR, Billac GB, Nichols DE, Nichols CD. 5-HT2A receptors: Pharmacology and functional selectivity. Pharmacol Rev. 2025 Jul 1;77(4):100059. doi:10.1016/j.pharmr.2025.100059

53. Halberstadt AL, Chatha M, Klein AK, Wallach J, Brandt SD. Correlation between the potency of hallucinogens in the mouse head-twitch response assay and their behavioral and subjective effects in other species. Neuropharmacology. 2020 May 1;167:107933. doi:10.1016/j.neuropharm.2019.107933 PubMed PMID: 31917152; PubMed Central PMCID: PMC9191653.

54. Andolina D, Puglisi-Allegra S, Ventura R. Strain-dependent differences in corticolimbic processing of aversive or rewarding stimuli. Front Syst Neurosci. 2015 Feb 4;8:207. doi:10.3389/fnsys.2014.00207 PubMed PMID: 25698940; PubMed Central PMCID: PMC4316691.

55. Belzung C, Griebel G. Measuring normal and pathological anxiety-like behaviour in mice: a review. Behav Brain Res. 2001 Nov 8;125(1):141–9. doi:10.1016/S0166-4328(01)00291-1

56. Crabbe JC, Wahlsten D, Dudek BC. Genetics of Mouse Behavior: Interactions with Laboratory Environment. Science. 1999 Jun 4;284(5420):1670–2. doi:10.1126/science.284.5420.1670

57. Woody ML, Gibb BE. Integrating NIMH Research Domain Criteria (RDoC) into depression research. Curr Opin Psychol. 2015 Aug 1;Depression4:6–12. doi:10.1016/j.copsyc.2015.01.004

