## Supplemental Material for "Identifying factors that contribute to the behavioral effects of psilocybin in preclinical mouse models"

#### Statistical Analysis

Coefficient of variation was calculated for head twitch, locomotion, SPT, and EPM by dividing the standard deviation (σ) by the mean (μ), and then multiplied by 100. NSF latency data was excluded from this analysis because the data is highly right-skewed and censored.

*Principal component analysis.* To examine the latent structure and assess the shared and distinct structure of behavioral outcomes across domains, we performed a principal component analysis (PCA) on SPT, EPM, and NSF risk. Only animals with complete data for all three behavioral measures were included in this analysis (N=464). Variables were centered and scaled prior to analysis, and PCA was conducted using a correlation-based approach. The NSF risk score was sign-flipped so that higher values consistently reflected better behavioral performance (i.e., lower anxiety) to facilitate interpretation of component loadings across domains. Pairwise Pearson correlations between behavioral measures were first examined. PCA was conducted on centered and scaled variables using singular value decomposition as implemented in the "prcomp" function in R. Component retention was determined using Kaiser's criterion (eigenvalue > 1) and proportion of variance explained. Component loadings indicate the contribution of each behavioral measure to the principal components, with values > |0.3| considered meaningful. Standardized loadings, eigenvalues, and variance explained are reported for all components. Communalities (sum of squared loadings) indicate the proportion of each variable's variance captured by the retained components. Principal component scores were extracted for each animal and used in subsequent analyses to test whether experimental factors (drug treatment, sex, age, strain, stress exposure, 5-HT1B expression) were associated with variance along the dominant behavioral dimensions. Linear regression models were used to assess main effects of these predictors on PC1, which captured the largest proportion of shared variance across behavioral measures.

*Multiple regressions and MANOVA*. To assess moderators of psilocybin's acute behavioral effects, we first conducted a univariate linear regression model for each acute behavioral outcome separately to maximize sample size. Models included drug treatment, age (mean-centered), sex, strain, stress, and 5-HT1BR, along with interaction terms between drug and each covariate to test moderation effects. Type III sums of squares were used to evaluate main effects and interactions. A reduced model excluding interactions was fit to assess robustness of main drug effects for locomotion too. To assess whether moderators altered the combined acute behavioral profile induced by psilocybin, a multivariate analysis of variance (MANOVA) was conducted on the subset of animals with both head-twitch and locomotor data available. Head-twitch and locomotion were treated as joint dependent variables in a multivariate linear model including the same predictors and interaction terms as the univariate analyses. Type III multivariate tests were performed using Pillai's trace All analyses were conducted in R using the "*car*" (Fox, J Weisrberg 2019) and "*emmeans*" (Lenth R 2025) packages. Distributions of sex, age, genotype, stress model, and treatment across behavioral measures are included in supplementary table 4.

*Structural equation modeling*. To assess which baseline predictors (sex, age, strain, stress 5-HT1Br) moderate the behavioral effects of psilocybin, we fit a multivariate regression model using structural equation modeling (SEM) using the lavaan package (v.0.6-20) in R. Sucrose preference (SPT) and elevated plus maze performance (EPM) were modeled simultaneously. Each outcome was regressed on drug condition (psilocybin vs. saline), age (mean centered), sex, 5-HT1B expression, strain (C57 vs. 129/C57), and stress exposure (no stress, corticosterone, repeated forced swim), as well as interaction terms between drug condition and each covariate to test for moderation effects. Residual correlations were freely estimated to account for shared variance between behavioral domains. Models were estimated using maximum likelihood with robust standard errors (MLR) to account for potential non-normality. Missing data were handled using full information maximum likelihood (FIML), allowing all available observations to contribute to parameter estimation under a missing-at-random assumption. Standardized coefficients (β), standard error, z-values, p-values, model fit indices including Tucker-Lewis index (TLI) and mean square root of approximation (RMSEA), and outcome-specific $R^2$ values were reported. Continuous predictors were normalized prior to inclusion to improve interpretability of interaction terms.

*Cox proportional hazard regression*. Novelty-suppressed feeding (NSF) score was analyzed using Cox proportional hazards regression to account for right-censored latency-to-feed data. Time to initiate feeding was modeled as a function of drug treatment, age, sex, 5-HT1B expression, strain, and stress condition. To test whether baseline predictors moderated drug effects, interaction terms between drug and covariates, as well as three-way interactions between drug, genotype, and stress, were included in the model. Animals that did not initiate feeding during the testing period were treated as censored observations. Hazard ratios (HRs) and corresponding confidence intervals were reported, with higher hazard indicating greater likelihood of feeding. Model assumptions were assessed by examining Schoenfeld residuals for proportional hazard violations. The global test indicated no significant departure from proportional hazards ($p>0.05$) and model concordance ($C = 0.671$) indicated overall predictive accuracy. In addition to survival modeling, a continuous NSF-derived anxiety index, nsf_risk, was computed to enable inclusion of NSF-related information in future principal component and machine learning analyses. Specifically, a null Cox proportional hazards model was fit to NSF latency data across animals. Deviance residuals were extracted and standardized to yield a z-scored risk index. This approach preserves information from both feeding and non-feeding animals while accounting for censoring, producing a continuous measure of anxiety-like behavior independent of experimental predictors.

*Random Forest Classification***.** A random forest classifier was trained to assess whether a multivariate signature of psilocybin treatment could be detected from biological and behavioral factors in 5-HT1BR-intact animals (n = 288). Predictors included sex, age, strain, stress exposure, sucrose preference (SPT), elevated plus maze performance (EPM), and NSF risk. Data were partitioned into a training set (70%) and a fixed held-out test set (30%) using stratified sampling to preserve class balance. To obtain stable estimates of model performance and variable importance, the random forest was refit 2,000 times on bootstrap resamples of the training data (sampling with replacement); the held-out test set remained fixed across all iterations. Each forest used 500 trees with mtry = $\sqrt{p}$. Model performance was summarized as the mean AUC across all 2,000 iterations with 95% confidence intervals derived from the bootstrap distribution. To assess whether classification performance exceeded chance, drug labels were randomly shuffled and the model was refit on a new stratified 70/30 split for each of 5,000 permutation iterations, generating a null AUC distribution; a one-sided p-value reflects the proportion of null AUCs exceeding the observed mean AUC. All analyses used the randomForest and pROC packages in R (seed = 123). Two additional models were run to assess bias and randomization of groups in drug assignment. One included baseline predictor only across the entire dataset, the other baseline predictors only within the subset of mice with 5-HT1BR intact signaling.

### Supplemental Results

#### The acute behavioral responses to psilocybin are influenced by the strain and age of mice

We first examined the effects of moderators on the psilocybin induced head twitching and locomotion individually. As expected, psilocybin induced a robust increase in head-twitch responding (main effect of drug: $F(1, 52) = 204.21$, $p < 0.0001$). This effect was significantly moderated by strain, such that C57BL/6J mice exhibited about 10 more head twitches compared to the mixed background 129;C57BL/6J mice within the 15-minute period after drug administration (Fig S1A; drug x strain interaction: $F(1, 52) = 81.18$, $\beta = -10.61 \pm 1.18$, $p < 0.0001$). However, C57BL/6J mice were treated with a concentration of 3mg/kg of psilocybin while 129;C57BL/6J with 5mg/kg so we cannot exclude a dose-dependent effects. Age also significantly moderated the acute head twitch response to psilocybin (drug x age interaction: $F(1, 52) = 6.41$, $p = 0.014$), with older animals exhibiting reduced head twitch responding to psilocybin (Fig. S1B). There was no significant moderation of psilocybin's effects on head twitching by sex or stress. Next, we examined locomotor activity in the first 30 minutes following drug administration. Psilocybin significantly decreased locomotion compared to saline (main effect of drug: $F(1, 78) = 15.79$, $p = 0.0002$), however there were no significant effects of moderators on the hypolocomotor response, suggesting psilocybin's response on movement is not impacted by age, sex, stress, or strain of the mouse.

#### MANOVA on head twitch and locomotion

Given that head twitch response and locomotion represent dissociable but related acute behavioral responses to psilocybin, we next performed a multivariate analysis of variance (MANOVA) on a subset of animals with both measures available (N=41) to assess moderation of the overall acute behavioral profile. This revealed a significant moderation of psilocybin's acute behavioral response by both mouse strain and stress paradigm (main effect of psilocybin: Pillai's trace = 0.77, $F(2, 28) = 47.05$, $p < 0.0001$; strain x psilocybin: Pillai's trace = 0.29, $F(2, 28) = 5.84$, $p = 0.008$; stress x psilocybin: Pillai's trace = 0.20, $F(2, 28) = 3.43$, $p = 0.047$)(Supp. S2). These results indicate that strain and stress alter the pattern of acute psilocybin-induced behaviors across an acute behavioral profile.

#### Random forest classification of baseline predictors only.

To confirm adequate randomization, a random forest classifier trained on baseline predictors only (age, sex, strain, stress; N = 469 5-HTBR signaling controls) achieved chance-level discrimination between drug groups (AUC = 0.526, permutation = 0.56). This indicates that saline and psilocybin groups were well-matched on biological and experimental variables prior to treatment. We also ran a random forest classifier trained on baseline predictors across the entire dataset for similar purposes (age, sex, strain, stress, 5-HT1BR; N = 693. The classifier also achieved chance-level discrimination between drug groups (AUC = 0.524, permutation = 0.52), indicating saline and psilocybin assignments were well randomized across biological and experimental variables.

**Supplemental Figures**

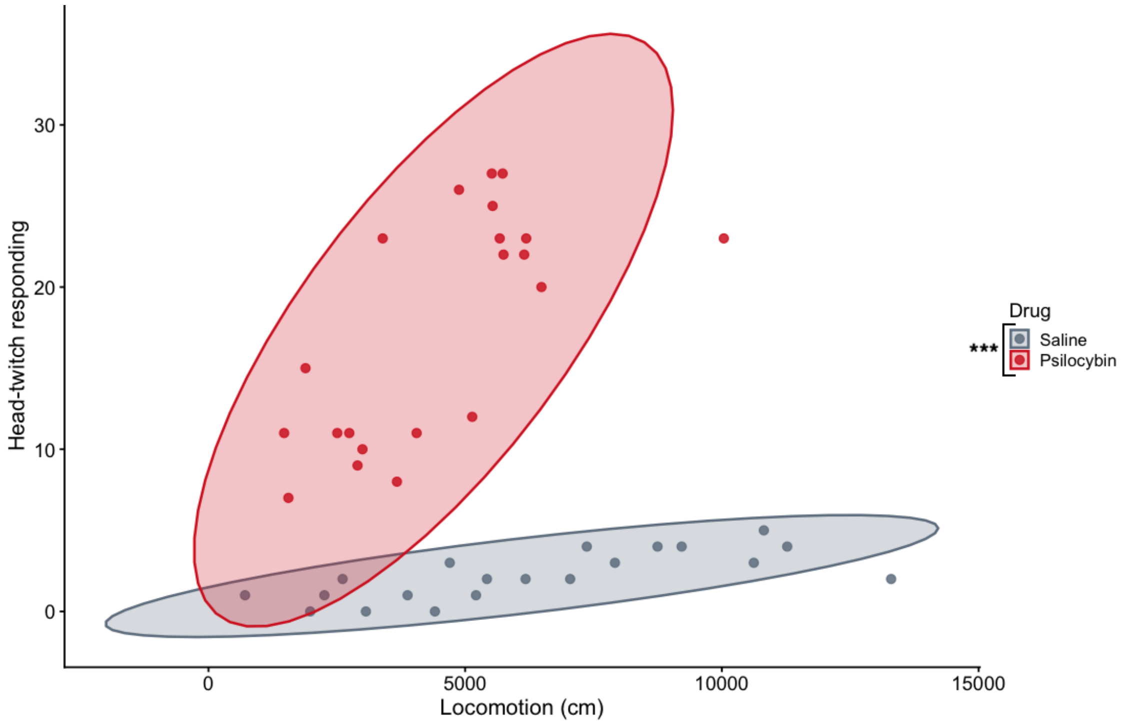


**Supplemental Figure S1**. **Psilocybin shifts the acute behavioral profile in mice.** A MANOVA shows that psilocybin significantly alters locomotion and head-twitching. The scatterplot shows individual animals with head-twitch responding plotted against locomotor activity measured during the acute drug period. Individual points represent individual mice colored by drug treatment (saline: grey, psilocybin: red). Ellipses represent 95% confidence regions assuming a bivariate t distribution for each treatment group. *** $p < 0.0001$

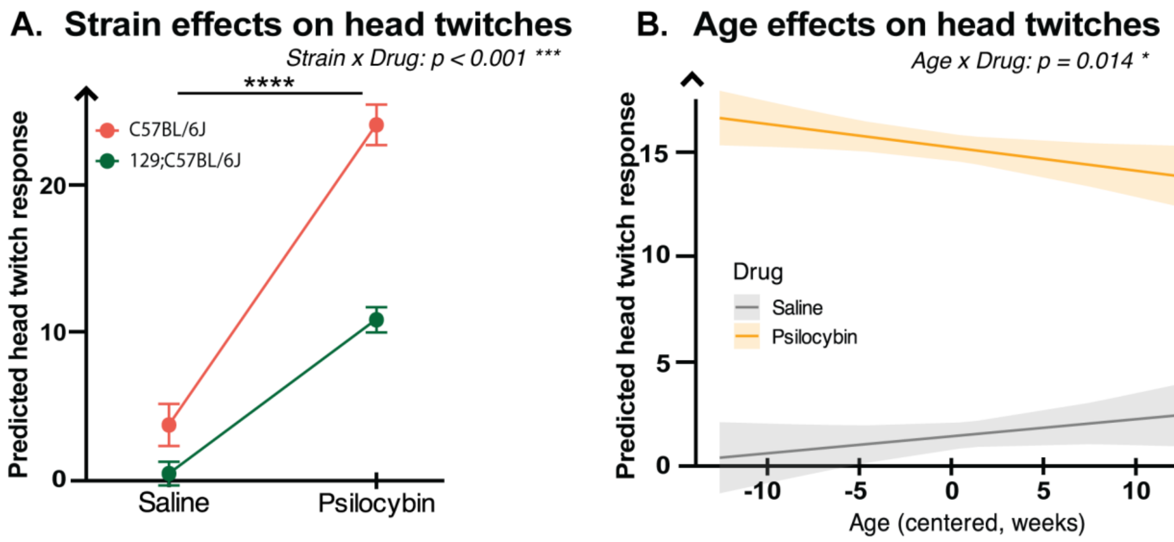


**Supplemental Figure S2. Strain and age moderate the head twitch response to psilocybin.** (A) Head twitch response following saline or psilocybin administration across mouse strains. Psilocybin significantly increases the number of head twitches in both the C57BL/6J (red) and the mixed 129S6/SvEv;C57BL/6J (green) strain. This effect was significantly greater in the C57BL/6J strain compared to 129S6/SvEv;C57BL/6J (strain x drug interaction $p < 0.001$). Points represent model-predicted means ± SEM. (B) Age-dependent effects of psilocybin on head-twitch response. Predicted head-twitch responses are plotted as a function of age (centered, weeks) for saline- and psilocybin-treated mice. Psilocybin-evoked head twitching declines with increasing age with no changes in head twitches across saline-treated mice (age x drug interaction $p = 0.014$). Lines represent model-predicted values with shaded 95% confidence intervals. Statistical significance is indicated as $p < 0.05$*, $p<0.001$***.

### Supplemental Tables

**Distribution of animals across behavioral outcomes**

| Behavioral assay | n | n_female | n_male | n_saline | n_psi | n_ctrl | n_ko | n_C57 | n_129/ C57 | n_unstressed | n_CORT | n_RFS | mean_age | sd_age | min_age | max_age |
|---|---|---|---|---|---|---|---|---|---|---|---|---|---|---|---|---|
| **HTR** | 64 | 23 | 41 | 32 | 32 | 47 | 17 | 21 | 43 | 25 | 27 | 12 | 22.6 | 7.2 | 10 | 36 |
| **Locomotion** | 85 | 35 | 50 | 46 | 39 | 67 | 18 | 33 | 52 | 41 | 32 | 12 | 28.6 | 8.1 | 10 | 43 |
| **SPT** | 562 | 247 | 315 | 293 | 269 | 379 | 183 | 126 | 436 | 156 | 296 | 110 | 23.5 | 8.7 | 10 | 46 |
| **EPM** | 594 | 276 | 318 | 308 | 286 | 388 | 206 | 76 | 518 | 135 | 292 | 167 | 23.9 | 9.5 | 10 | 48 |
| **NSF** | 554 | 237 | 317 | 291 | 263 | 369 | 185 | 128 | 426 | 162 | 295 | 97 | 23.7 | 9.1 | 10 | 46 |

**Supplemental Table 1. Distribution of animals across behavioral measures.** Total numbers of animals run in each behavioral assay is shown (n). Ns are also shown stratified by sex (female, male), drug condition (sal, psi), 5-HT1BR expression (cntrl, ko), strain (C57, mixed 129/C57), and stress paradigm (unstressed, CORT, or RFS). Mean, standard deviation, min, and max ages reported in weeks are presented in the last 4 columns.

| Predictor | β (Estimate) | SE | t value | p value |
|---|---|---|---|---|
| Intercept | 0.561 | 0.159 | 3.53 | **0.0005** *** |
| Drug (Psil vs Sal) | 0.263 | 0.099 | 2.66 | **0.008** ** |
| Sex (M vs. F) | -0.324 | 0.106 | -3.07 | **0.002** ** |
| 5-HT1BR | 0.688 | 0.102 | 6.75 | **< 1e-10** *** |
| Age | -0.026 | 0.007 | -3.97 | **< 0.001** *** |
| Strain | -0.171 | 0.139 | -1.23 | 0.219 |
| CORT_stress | -0.981 | 0.127 | -7.71 | **< 1e-13** *** |
| RFS_stress | -0.594 | 0.155 | -3.84 | **< 0.001** *** |

***Model fit:*** *$R^2 = 0.195$, Adjusted $R^2 = 0.182$; $F(7, 456) = 15.74$, $p < 2 \times 10^{-16}$*

**Supplemental Table 2**. **Linear regression predicting PC1 score.** PC1 was derived from principal component analysis of SPT, EPM, and NSF measures. Psilocybin treatment, females, 5-HT1BR– mice, younger mice, and unstressed mice are associated with higher PC1 scores than saline-treated mice, males, intact 5-HT1BR signaling, older mice, and mice that underwent CORT stress or RFS stress. Strain had no significant effect on PC1 predictions. Positive coefficients indicate higher PC1 scores, meaning a higher positive affective behavioral phenotype. ** p < 0.001, ** p < 0.01

| Predictor | SPT | | | EPM | | |
|---|---|---|---|---|---|---|
| | β (SE) | z | p | β (SE) | z | p |
| **Main effects** | | | | | | |
| Age | −0.113 (0.036) | -3.14 | **0.002** ** | −0.389 (0.289) | -1.34 | 0.179 |
| Sex | −1.422 (0.556) | -2.56 | **0.011** * | 9.354 (4.438) | 2.11 | **0.035** * |
| 5-HT1BR | 0.963 (0.646) | 1.49 | 0.136 | 11.320 (4.338) | 2.61 | **0.009** ** |
| Strain | 0.949 (0.570) | 1.67 | 0.096 | −37.429 (9.550) | -3.92 | **<0.001** *** |
| Drug | 2.352 (1.051) | 2.24 | **0.025** * | 27.238 (15.200) | 1.79 | 0.073 |
| CORT_stress | −2.374 (0.651) | -3.65 | **<0.001** *** | −30.077 (6.457) | -4.66 | **<0.001** *** |
| RFS_stress | −1.973 (0.789) | -2.50 | **0.012** * | −32.886 (5.133) | -6.41 | **<0.001** *** |
| **Drug × Moderator interactions** | | | | | | |
| Drug × Age | −0.007 (0.065) | -0.11 | 0.913 | −0.450 (0.368) | -1.22 | 0.221 |
| Drug × Sex | −2.077 (0.842) | -2.47 | **0.014** * | −19.215 (6.249) | -3.08 | **0.002** ** |
| Drug × 5-HT1BR | 0.244 (0.978) | 0.25 | 0.803 | −1.609 (5.922) | -0.27 | 0.786 |
| Drug × Strain | 0.547 (0.863) | 0.63 | 0.527 | 8.659 (13.361) | 0.65 | 0.517 |
| Drug × CORT | −0.994 (0.942) | -1.06 | 0.291 | −31.493 (9.987) | -3.15 | **0.002** ** |
| Drug × RFS | 0.374 (1.276) | 0.29 | 0.769 | −17.131 (9.024) | -1.90 | 0.058 † |

Significance: *** $p < 0.001$, ** $p < 0.01$, * $p < 0.05$, † $p < 0.10$

*Model fit: SPT $R^2 = 0.142$, EPM $R^2 = 0.358$*

**Supplemental Table 3. Multivariate regression of sucrose preference score and elevated plus maze.** Structural equation modeling including interaction terms was performed to simultaneously estimate predictors of SPT and EPM while accounting for residual covariance. There was a main effect of age (p = 0.002), sex (p = 0.011), drug (p = 0.025), and both stress models (CORT: p < 0.001, RFS: p = 0.012) on SPT. There was a main effect of sex (p = 0.035), 5-HT1BR (p= 0.009), strain (p < 0.001), and stress models (p < 0.001) in the EPM. Drug effects were moderated by sex in both the SPT (p = 0.014) and EPM (p = 0.002). It was also moderated by stress, especially CORT stress (p = 0.002), in the EPM.

| Predictor | β (log-HR) | SE | z | p | HR (expβ) | 95% CI HR |
|---|---|---|---|---|---|---|
| Age | -0.0037 | 0.0095 | -0.39 | 0.697 | 0.996 | [0.978, 1.015] |
| Strain | 0.028 | 0.184 | 0.15 | 0.878 | 1.03 | [0.718, 1.474] |
| CORT_stress | **-0.563** | 0.179 | -3.14 | **0.0017** | **0.57** | **[0.40, 0.81]** |
| RFS_stress | 0.214 | 0.201 | 1.07 | 0.286 | 1.24 | [0.84, 1.84] |
| Sex | 0.155 | 0.306 | 0.51 | 0.612 | 1.17 | [0.64, 2.13] |
| Psilocybin | **0.775** | 0.306 | 2.53 | **0.011** | **2.17** | **[1.19, 3.96]** |
| 5-HT1BR | **1.319** | 0.312 | 4.24 | **<0.001** | **3.74** | **[2.03, 6.89]** |
| Sex × Drug | -0.758 | 0.396 | -1.91 | 0.056 | 0.47 | [0.22, 1.02] |
| Sex × 5-HT1BR | 0.109 | 0.398 | 0.27 | 0.784 | 1.12 | [0.51, 2.43] |
| Drug × 5-HT1BR | **-1.007** | 0.421 | -2.39 | **0.017** | **0.37** | **[0.16, 0.83]** |
| Sex × Drug × 5-HT1BR | **1.147** | 0.553 | 2.08 | **0.038** | **3.15** | **[1.07, 9.30]** |

*Model fit: concordance = 0.676; likelihood ratio test $\chi^2(11) = 90.6$, $p < 1\times10^{-14}$; N = 554 animals.*

**Supplemental Table 4. NSF Cox model with three-way interaction of sex by drug by 5-HT1BR.** Cox regression was performed to account for right-censoring at the test cutoff. This 3-way model included age, strain, stress models, sex, drug, 5-HT1BR, all 2-way interactions of drug, sex, and 5-HT1BR, and the 3-way interaction between sex, drug, and 5-HT1BR. Psilocybin and 5-HT1BR– was associated with increased hazard of feeding (shorter latencies to eat), while CORT stress significantly reduced feeding hazard. A significant drug x 5-HT1BR (p = 0.017) and trend level drug x sex (p = 0.056) interactions were detected. There was a significant sex x drug x 5-HT1BR (p = 0.038), indicating that the effect of psilocybin on feeding latency is dependent on 5-HT1BR signaling and sex.

**References:**


Fox J, Weisberg S (2019). _An R Companion to Applied Regression_, Third edition. Sage, Thousand Oaks CA. <https://www.john-fox.ca/Companion/>.

Lenth R (2025). _emmeans: Estimated Marginal Means, aka Least-Squares Means_. doi:10.32614/CRAN.package.emmeans <https://doi.org/10.32614/CRAN.package.emmeans>, R package version 1.11.2-8, <https://CRAN.R-project.org/package=emmeans>.
